# Interspecies Social Spreading: Interaction between two sessile soil bacteria leads to emergence of surface motility

**DOI:** 10.1101/296814

**Authors:** Lucy M. McCully, Adam S. Bitzer, Sarah C. Seaton, Leah M. Smith, Mark W. Silby

## Abstract

Bacteria often live in complex communities in which they interact with other organisms. Consideration of the social environment of bacteria can reveal emergent traits and behaviors that would be overlooked by studying bacteria in isolation. Here we characterize a social trait which emerges upon interaction between the distantly-related soil bacteria *Pseudomonas fluorescens* Pf0-1 and *Pedobacter* sp. V48. On hard agar, which is not permissive for motility of the mono-culture of either species, co-culture reveals an emergent phenotype we term ‘interspecies social spreading,’ where the mixed colony spreads across the hard surface. We show that initiation of social spreading requires close association between the two species of bacteria. Both species remain associated throughout the spreading colony, with reproducible and non-homogenous patterns of distribution. The nutritional environment influences social spreading; no social behavior is observed under high nutrient conditions, but low nutrient conditions are insufficient to promote social spreading without high salt concentrations. This simple two-species consortium is a tractable model system that will facilitate mechanistic investigations of interspecies interactions and provide insight into emergent properties of interacting species. These studies will contribute to the broader knowledge of how bacterial interactions influence the functions of communities they inhabit.

**Importance:** The wealth of studies on microbial communities has revealed the complexity and dynamics of the composition of communities in many ecological settings. Fewer studies probe the functional interactions of the community members. Function of the community as a whole may not be fully revealed by characterizing the individuals. In our two-species model community, we find an emergent trait resulting from the interaction of the soil bacteria *Pseudomonas fluorescens* Pf0-1 and *Pedobacter* sp. V48. Observation of emergent traits suggests there may be many functions of a community that are not predicted based on *a priori* knowledge of the community members. These types of studies will provide a more holistic understanding of microbial communities, allowing us to connect information about community composition with behaviors determined by interspecific interactions. These studies increase our ability to understand communities, such as the soil microbiome, plant-root microbiome, and human gut microbiome, with the final goal of being able to manipulate and rationally improve these communities.

## Introduction

Within soils live a plethora of microbial species that form complex communities responsible for important ecological functions, such as nutrient cycling and plant health. Omics approaches have given us a wealth of information on the composition, diversity, metabolic potential, and ecology of plant- and soil-associated microbial communities (1, 2). However, to get a complete understanding of microbial functions and interactions within these environments, we must look at every layer, from the full community *in vivo* to the individual microbe *in vitro* (3). Historically, research has focused on the study of single species in pure culture, but bacteria are social organisms. Thus, the study of the mechanisms and consequences of multi-species interactions is necessary for us to understand the function of microbial communities as a whole. Investigating entire soil communities *in situ* presents considerable challenges because of fluctuating soil conditions and the wide range of relevant scales, ranging from particulate to ecological levels (2). Reducing the microbial community to pair-wise interactions or small consortia allows for a detailed mechanistic study. This reduction is also an essential link between studying isolated microbes in the laboratory and understanding the collective activities of natural microbial communities (4).

Recent work has considered the social environment of bacteria, investigating altered behaviors and production of secondary metabolites when co-cultured with other organisms. Some bacteria exhibit emergent behaviors when presented with other species, likely the result of induction of genes that are not expressed in pure culture. For example, some *Pseudomonas fluorescens* strains produce an antifungal compound during interactions with other species (5–9). The co-culture of different actinomycete species results in the production of secondary metabolites, changes in pigment, and sporulation (10–12). The presence of *E. coli* or *Pseudomonas* species effects sporulation and biofilm formation in *Bacillus subtilis* (13, 14). One subset of social interactions are those which alter the motility behaviors and capabilities of other species. For example, physical association with *Saccharomyces cerevisiae* results in *Streptomyces venezuelae* consuming the yeast and triggers ‘exploratory growth’ of the bacteria (15). This exploration is not observed when S. *venezuelae* is grown in mono-culture, under the same environmental conditions. In another example, *B. subtilis* moves away from a *Streptomyces* competitor across a solid surface, but does not do so in isolation (16, 17). Other behaviors appear less competitive, where a motile species will travel with a non-motile species that can degrade antibiotics, allowing the consortium to colonize hostile environments (18, 19). *Xanthomonas perforans* can even change the behavior of *Paenibacillus vortex*, producing a signal that induces *P. vortex* to swarm towards it so it can hitchhike (20).

*Pseudomonas fluorescens* Pf0-1 and *Pedobacter* sp. V48 are known to interact though diffusible and volatile signals, which induce production of an antifungal compound by *P. fluorescens* (6–8). Previous studies with *Pedobacter* and a strain closely-related to *P. fluorescens* Pf0-1 (AD21) found that the mixture of the strains showed reciprocal gene expression changes and antagonistic behavior toward the plant pathogen *Rhizoctonia solani* (5, 9). The initial study noted expansion of the mixed strains beyond the initial area of inoculation (5), but the phenotype was not characterized, and has not been the focus of any further studies. We investigated this observation using a new assay. Instead of culturing *P. fluorescens* Pf0-1 and *Pedobacter* without contact, as was done in the antagonism assays (6), we mix them together. We hypothesized that, while antibiotic production can be induced at a distance through diffusible or volatile signals, the motility behavior requires close contact and is therefore controlled in a manner distinct from the other two forms of communication.

In this study, we describe an interaction between two distantly-related soil bacteria, *P. fluorescens* Pf0-1 (phylum: Proteobacteria) and *Pedobacter* sp. V48 (phylum: Bacteroidetes). This interaction produces an emergent behavior, which we term “interspecies social spreading,” in which the bacteria move together across a hard agar surface. When grown in isolation, neither species moves beyond the typical amount of colony expansion. In co-culture, both bacteria are present throughout the spreading colony, and fluorescent imaging shows a non-homogenous distribution. We demonstrate that a close association between the colonies of both species is required for spreading to initiate and that the levels of nutrients and salts in the media affect the development of the spreading phenotype.

## Results

### Interspecies social spreading arises when mixing two distantly-related bacteria

In previous studies, antifungal activity was observed when *P. fluorescens* Pf0-1 and *Pedobacter* sp. V48 were cultured 15 mm apart (6). In addition to this interaction-induced trait, the possibility of motility was noted, but not further investigated, in a mixture of *P. fluorescens* AD21 and *Pedobacter* (5, 9). To explore this phenomenon, we developed an assay in which the induced motility is greater and more easily observed. Our approach differed from the conditions of the original observation in inoculation method, strain combination, and media composition. When we plated *P. fluorescens* Pf0-1 and *Pedobacter* on TSB-NK medium solidified with 2% agar a mixed colony of the two bacteria expanded across the surface of the agar, an environment in which neither monoculture exhibited motility. The emergent social spreading is shown in Fig. 1.

**Figure 1.**
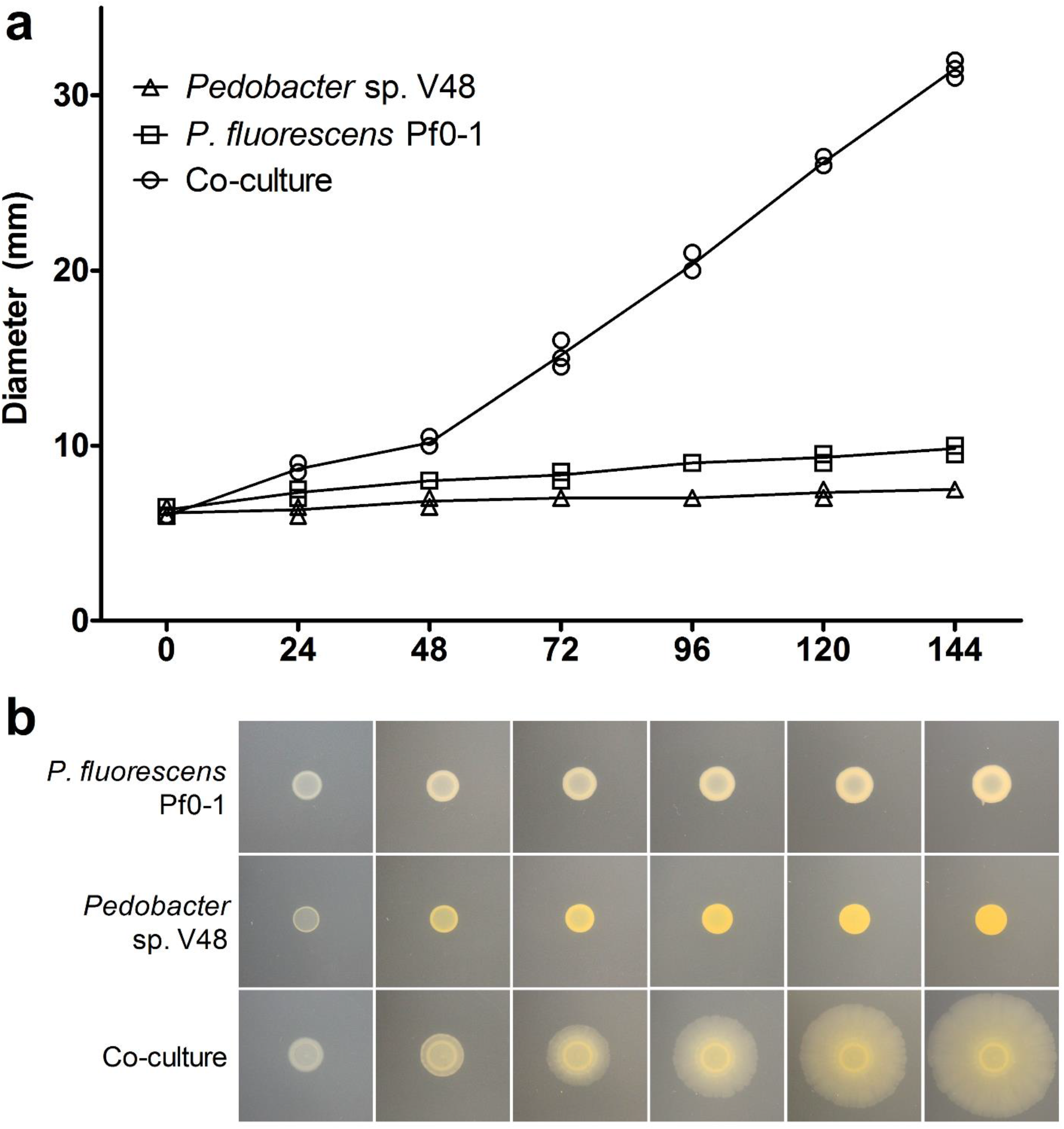
Mixed colony of *P. fluorescens* Pf0-1 and *Pedobacter* sp. V48 spreads across a hard surface (2% agar), a behavior not observed in the mono-culture of either species. a) Diameter of colonies at 24 h intervals for three independent experiments. b) Phenotypes of mono- and co-cultures at 24 h intervals. Contrast and brightness levels were adjusted for optimal viewing.

Social spreading becomes apparent between 24 and 48 h after inoculation, when the colony begins to spread from the edge of the inoculum (Fig. 1b 48 h). The diameter of the spreading co-culture is significantly different from the colony expansion of the mono-cultures starting at the 24 h time point (p < 0.001) (Fig. 1a). Once the spreading phenotype is fully visible (around 72 h), the average speed of expansion is 1.69 μm/min +/- 0.09 (SEM). At the onset of movement, the leading edge has a visibly thicker front (Fig. 1b 48 h). As the colony spreads, the thick front disappears and small ‘veins’ radiating from the center develop. Over time, the ‘veins’ become more pronounced towards the leading edge, making a ‘petal’ pattern (Figs. 2a, b). The leading edge is characterized by a distinctive, terraced appearance comprised of three to six layers (Fig. 2c). Varying the initial ratios between 5:1 and 1:5 *Pseudomonas:Pedobacter* did not have a visible effect on spreading across the plate (data not shown).

**Figure 2.**
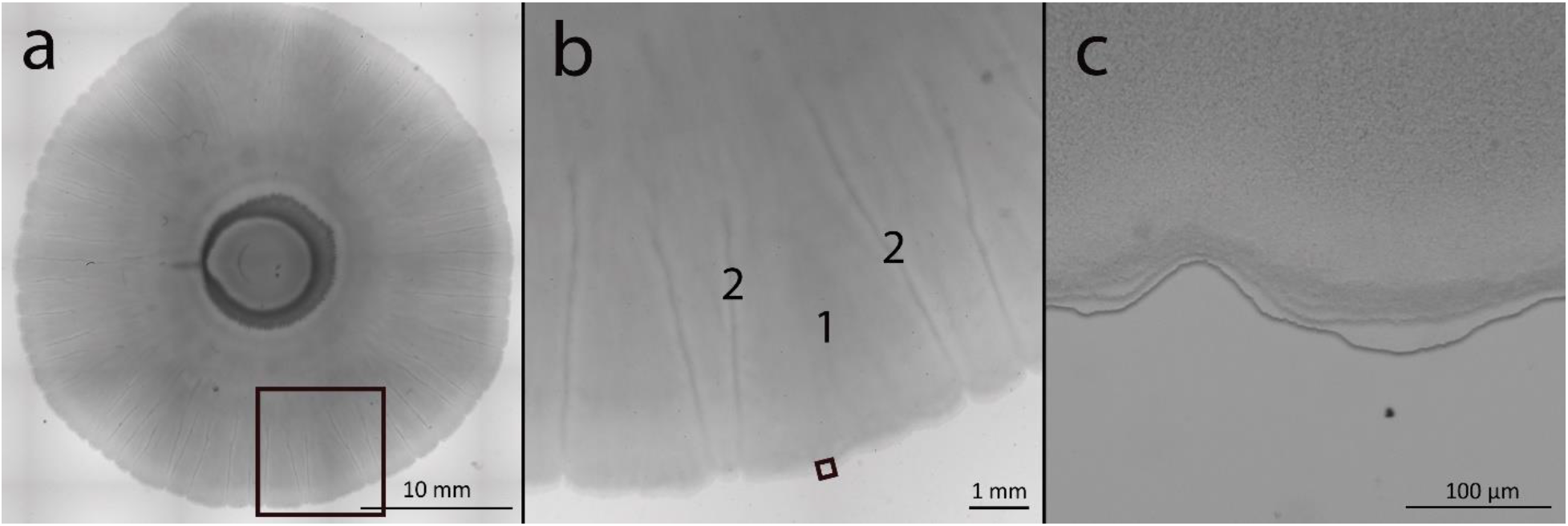
Mixed colony of *P. fluorescens* Pf0-1 and *Pedobacter* sp. V48 at different magnifications. a) Image of the whole co-culture colony created by stitching an 8X magnification mosaic. b) 16X magnification of the leading edge showing the patterns of ‘petals’ (1) in between ‘veins’ (2) visible near the edge of the colony c) 112X magnification shows a terraced appearance of the leading edge. Colony imaged 144 h after inoculation. Black boxes indicate area enlarged in the adjacent panel. Scale bars are noted at the bottom of each image. Contrast and brightness levels were adjusted for optimal viewing.

### *P. fluorescens* Pf0-1 and *Pedobacter* sp. V48 co-migrate

The previously observed ‘bacterial expansion’ of *Pedobacter* when interacting with *Pseudomonas* sp. AD21 was suggested to be gliding motility, triggered as a mechanism to escape competition from *P. fluorescens* (5, 9). We examined the possibility that the spreading observed when co-inoculating *Pedobacter* and *P. fluorescens* Pf0-1 was a result of *Pedobacter* moving away from *P. fluorescens*. Bacteria were collected from the center, middle, and edge of a seven-day-old motile colony. The presence or absence of each species was tested by culturing these samples on selective media. We recovered both species from each point in the spreading colony (data not shown), showing co-migration rather than an escape strategy by *Pedobacter*.

### Interspecies social spreading shows reproducible spatial organization

To obtain a more detailed look at the spatial relationships within the spreading colony, we tagged *P. fluorescens* with a cyan fluorescent protein (eCFP (21)) and *Pedobacter* with a red fluorescent protein (dsRedEXPRESS (21)), integrated into the chromosome. In *P. fluorescens*, eCFP carried by a miniTn7 transposon was integrated upstream of *glmS* (22), creating Pf0-ecfp. In *Pedobacter*, dsRedEXPRESS carried by the *HimarEm* transposon (23) was integrated at random locations in the chromosome, resulting in 16 independently-derived mutants with an insert. Each tagged *Pedobacter* strain (V48-dsRed) was indistinguishable from the wild-type in social assays with *P. fluorescens*, indicating no deleterious impact of the insertions. We picked one strain with an insert in locus N824_RS25465 (GenBank accession NZ_AWRU01000034), and no apparent defect in social spreading. The initiation of social spreading appeared slightly delayed in a mixture of the tagged strains, but the visible patterns and stages of development looked identical, and speed was not significantly different once spreading initiated.

Fluorescent microscopy verified culturing data that showed both bacteria are present throughout the spreading colony, but we also found that population density varies across distinct areas within the colony. These distribution patterns were highly reproducible and show six distinct zones (Fig. 3). At zone 1, the point of inoculation, fluorescent imaging shows a homogenous mix of both bacteria (Fig. 3b, c). Zone 2, the coffee ring effect formed at the edge of the point of inoculation (24–26), is bright orange, indicating that *Pedobacter* dominates this region (Fig. 3b). *Pedobacter* spreads out from this dense area into zone 3, in a starburst pattern (Fig. 3b). Just outward from the starburst, we see a blue ring (zone 4), where *P. fluorescens* appears more abundant (Figs. 3c, d). In the main body of the co-culture, a thin motile section spreads out, making ‘petals’ (Zone 5), with ‘veins’ (Zone 6) between them (Figs. 2b and 3a).

**Figure 3.**
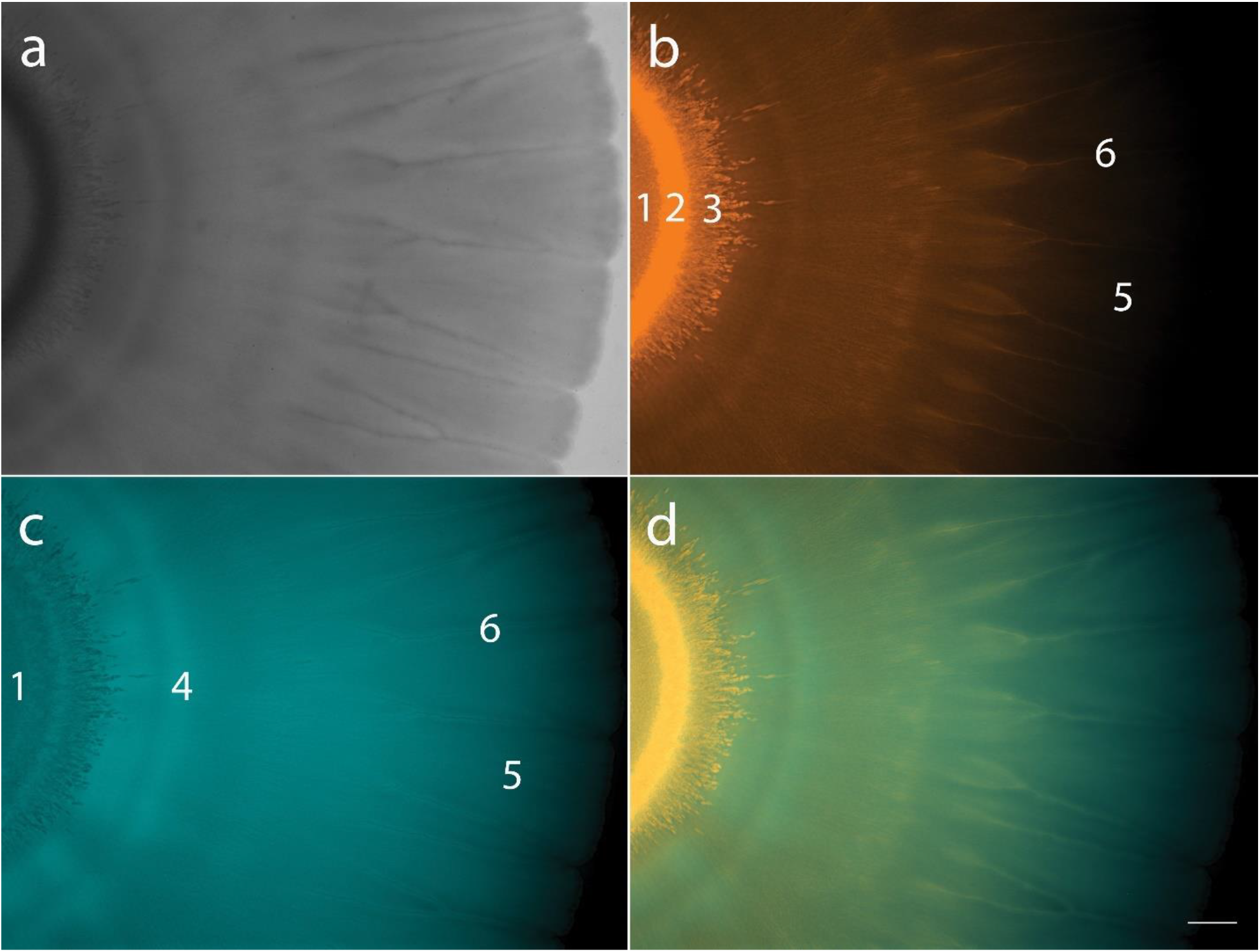
Mixed colony of fluorescently-tagged *P. fluorescens* Pf0-1 (Pf0-*ecfp*) and *Pedobacter* sp. V48 (V48-*dsRed*). a) Co-culture colony viewed with white light. b) Co-culture imaged using DsRed filter (filter set 43 HE), pseudo-colored in orange, showing *V48-dsRed* distribution throughout the colony. c) Co-culture imaged using CFP filter (filter set 47 HE), pseudo-colored in turquoise, showing Pf0-*ecfp* distribution throughout the colony. d) Merged images of DsRed and CFP filters. Numbers on panels b and c indicate six zones of distinct patterns: 1. Point of inoculation, 2. Coffee ring, 3. Starburst, 4. *P. fluorescens* ring, 5. Petals, 6. Veins. Colonies imaged at 7X magnification, scale bar represents 1 mm. Colony imaged 144 h after inoculation.

The ‘veins’ between the ‘petals’ appear to have high *Pedobacter* populations (Fig. 3b), while the areas directly surrounding them are dominated by *P. fluorescens* (Fig. 3c). The flat areas of the ‘petals’ appear more well-mixed, though the red signal becomes difficult to detect toward the edge of the colony (Fig. 3d). Overall, imaging data show that we can find both species throughout the colony, but the distribution is not homogenous. Rather, we observed reproducible patterns with some well-mixed areas and others of high spatial assortment.

### Diffused compounds and heat-killed cells do not trigger interspecies social spreading

Previous studies demonstrated that interactions between *P. fluorescens* and *Pedobacter* were mediated via both diffusible and volatile signals (6–8). We first asked whether spreading could be triggered by diffusible compounds produced by one of the partner species or by the co-culture. Mono and co-cultures grown on cellophane membranes were used to pre-condition our spreading assay agar. After two days, cellophane membranes (and the bacteria growing on them) were peeled off the agar. Plates were then inoculated with one of the partner strains to evaluate development of the social spreading phenotype. After seven days of growth, no sign of spreading beyond normal colony expansion was observed (change in diameter was not significantly different from negative control), indicating that no motility-inducing compounds had been secreted into the agar.

As the signal did not appear to diffuse through cellulose, we next asked if inactive cells or cell fragments of each species could trigger spreading in the other species. To address this, we used dead cells from one species, or from the spreading co-culture. Mono- and co-cultures were grown on TSB-NK media (as previously described) for 4 days, suspended in phosphate buffer, and heat-killed at 65 °C for 15 minutes. This heat-killed suspension was added directly on top of growing colonies of each species, or to wells adjacent to the colony being tested. Heat-killed suspension was added every 24 hours for five days. The plates were monitored for ten days, but no social spreading was observed under any condition, beyond that due to physical disruption which is also present in the buffer control.

### Physical association of *P. fluorescens* Pf0-1 and *Pedobacter* V48 is required for interspecies social spreading

Because a diffusible signal was unlikely to be triggering social spreading, we asked whether a close association between the two bacteria was a necessary condition for the interaction. To answer this question, we used assays in which the bacterial participants were plated side-by side with no physical barrier and in which they were separated only by semi-permeable membranes.

When colonies were adjacent, rather than mixed, no social spreading was observed while the *P. fluorescens* and *Pedobacter* colonies were visibly separate (data not shown). However, once the colonies grew sufficiently to make contact (Fig. 4 24 h), the colony started to spread out from the point of contact (72 h). The spreading front radiates outward (96 h), first developing around the *P. fluorescens* colony (144 h), then proceeding to surround the *Pedobacter* colony (192 h). At this level of resolution, contact between the colonies appears to occur before any social spreading can be seen.

**Figure 4.**
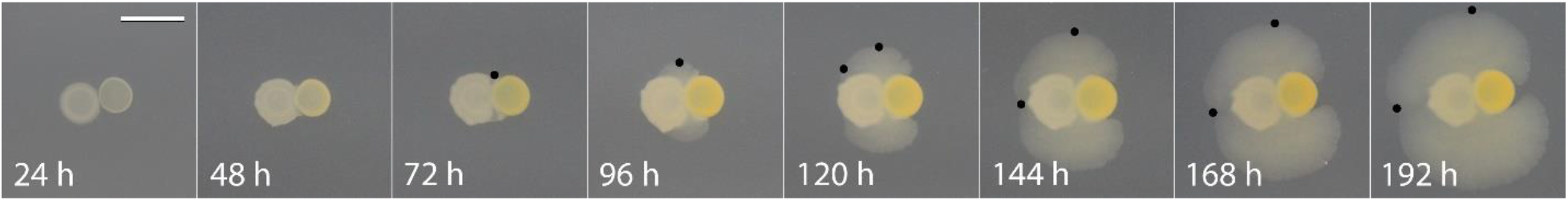
Social spreading emerges after contact between colonies of *P. fluorescens* Pf0-1 and *Pedobacter* sp. V48, left and right in each panel, respectively. Colonies come into contact 24 hours after inoculation; the motile front becomes visible 48 hours after contact and spreads outward and around the *P. fluorescens* colony before surrounding the *Pedobacter* colony. Black spots indicate sampling locations. Pictures taken every 24 h. Scale bar represents 10 mm.

Samples were collected from the edge of the moving front every 24 hours after contact, both on a y-axis from the point of contact and following the moving front as it wrapped around the *P. fluorescens* colony (Fig. 4). The presence of each species was tested by culturing these samples on selective media. Both species were culturable at every point sampled (data not shown), showing that *Pedobacter* is present in the moving front behind the *P. fluorescens* colony (Fig. 4, 144 h), on the opposite side of where they initially came into contact. This indicates that *Pedobacter* moves around the *P. fluorescens* colony on the motile front.

To further evaluate the requirement that *P. fluorescens* and *Pedobacter* be physically associated, we inoculated both strains immediately adjacent to each other but separated by either semi-permeable mixed-ester cellulose or PES (polyethersulfone) membranes. When inoculated this way, individual colony growth continued as normal, but these bacteria were unable to trigger social spreading despite their close proximity. After six days of growth, no sign of interspecies social spreading was observed (Fig. 5).

**Figure 5.**
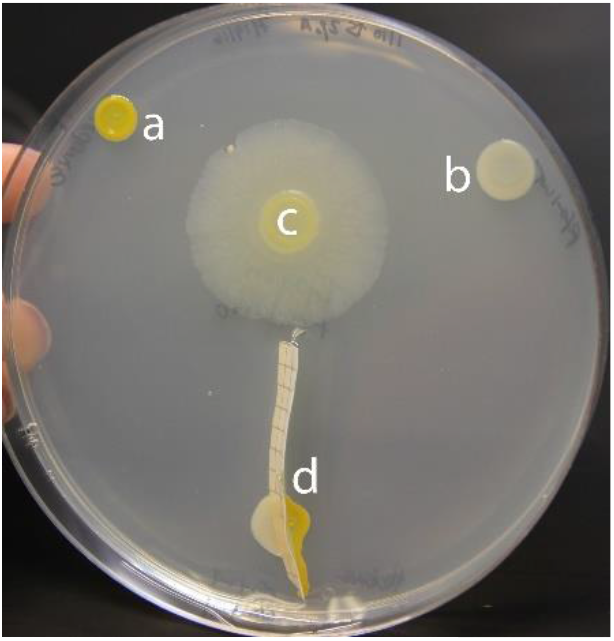
A semi-permeable barrier prevents development of the interspecies social spreading phenotype. a) *Pedobacter* sp. V48 monoculture, b) *P. fluorescens* Pf0-1 monoculture, c) a mixed colony, and d) *P. fluorescens* and *Pedobacter* separated by a mixed-ester cellulose membrane. Pictures taken 144 h after inoculation. Colonies were grown on a 100 mm petri dish.

### Nutritional environment influences interspecies social spreading

Conditions in soil and rhizosphere environments fluctuate, with bacteria subjected to a wide range of environmental stressors, including limited nutrient and water availability (2). Because such fluctuations may influence expression of traits, we examined the effect of nutrient level on interspecies social spreading. Our standard assay condition, TSB-NK, consists of 10% strength Tryptic Soy (3 g/L) supplemented with NaCl (5 g/L) and KH_2_PO_4_ (1 g/L).

We first asked if interspecies social spreading could initiate under richer nutrient conditions. No social spreading was apparent when *P. fluorescens* and *Pedobacter* were mixed on full-strength TSB (30 g/L) (Fig. 6a), with the co-culture exhibiting the same characteristics and colony expansion as the *P. fluorescens* mono-culture. We next asked whether the salt amendments to TSB-NK influence interspecies social spreading, using assays without the addition of salts, and with the addition of NaCl and KH_2_PO_4_ individually. When grown on 10% TSB, the co-culture is motile, but the distance spread is modest compared to when the medium is supplemented with both salts (Fig. 6c). The individual *P. fluorescens* colony expands similarly to the co-culture, suggesting minimal social behavior under these conditions. Growth on TSB-K changes neither pattern nor rate of mono- and co-culture expansion compared to 10% TSB (data not shown). On TSB-N, the mixed culture spreads and develops the patterns characteristic of interspecies social spreading, while the *P. fluorescens* mono-culture does not expand (Fig. 6d). The phenotype and diameter of the spreading colony are most similar to those observed in TSB-NK conditions (Fig. 6b).

**Figure 6.**
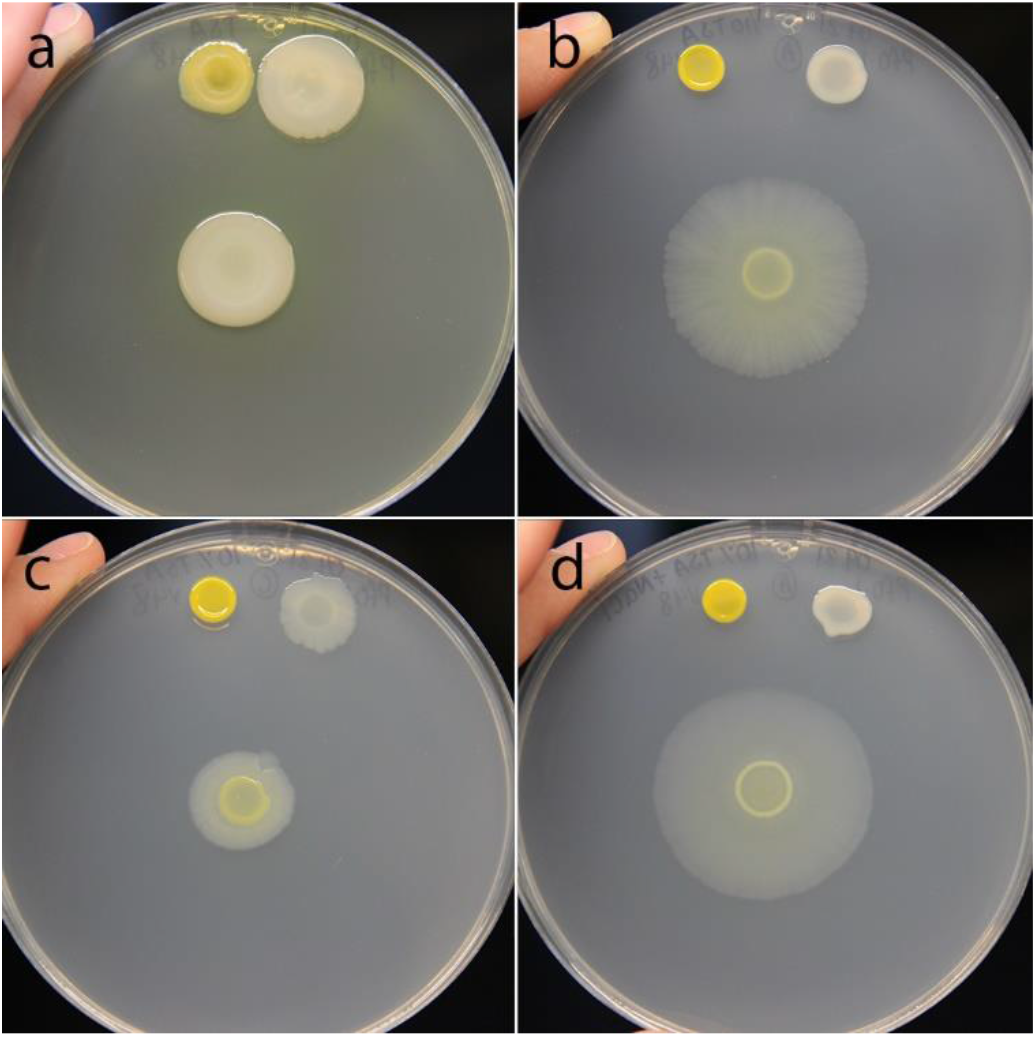
Low nutrient and high salt conditions are required for interspecies social spreading. a) Mixed colony on full-strength TSB does not show social spreading. b) Mixed colony on TSB-NK (10% Tryptic Soy supplemented with both NaCl and KH_2_PO_4_) shows social spreading. c) Mixed colony on 10% strength TSB shows impaired social spreading. d) Mixed colony on TSB-N (supplemented with NaCl) exhibits the interspecies social spreading phenotype. For all panels *Pedobacter* sp. V48 mono-culture is on the top left of the plate, *P. fluorescens* Pf0-1 is on the top right of the plate, and the mixed colony is in the center. Pictures were taken 144 h after inoculation. Colonies were grown on a 100 mm petri dish.

In the previous experiment, we observed that variations of Tryptic Soy media led to altered social phenotypes. To assess the influence of each component of TSB on interspecies social spreading, we utilized a medium in which these were individually manipulated. We made eight combinations of media to vary D-glucose, tryptone, and NaCl in concentrations equivalent to those in full-strength and 10% TSB. On media with D-glucose or tryptone at full-strength concentrations, we did not observe social spreading regardless of the concentration of the other components (Figs. 7a-f). In these conditions, the appearance and expansion of the co-culture resembled that of the *P. fluorescens* mono-culture, with notably greater biomass in media with full-strength tryptone (Figs. 7a-d). When the concentration of all three components was reduced to 10% we observed social spreading, but the migration distance of the co-culture was modest, and *P. fluorescens* mono-culture expanded to a similar extent (Fig. 7h). On media containing 10% strength D-glucose, 10% strength tryptone, and full-strength NaCl, interspecies social spreading emerged when *P. fluorescens* and *Pedobacter* were co-cultured (Fig. 7g). Unique to this condition, the mono-cultures of both strains are immotile, indicating a dramatic change in behavior when strains are mixed. The observations under this condition are most similar to those observed on TSB-N and TSB-NK (Figs. 6b, d).

**Figure 7.**
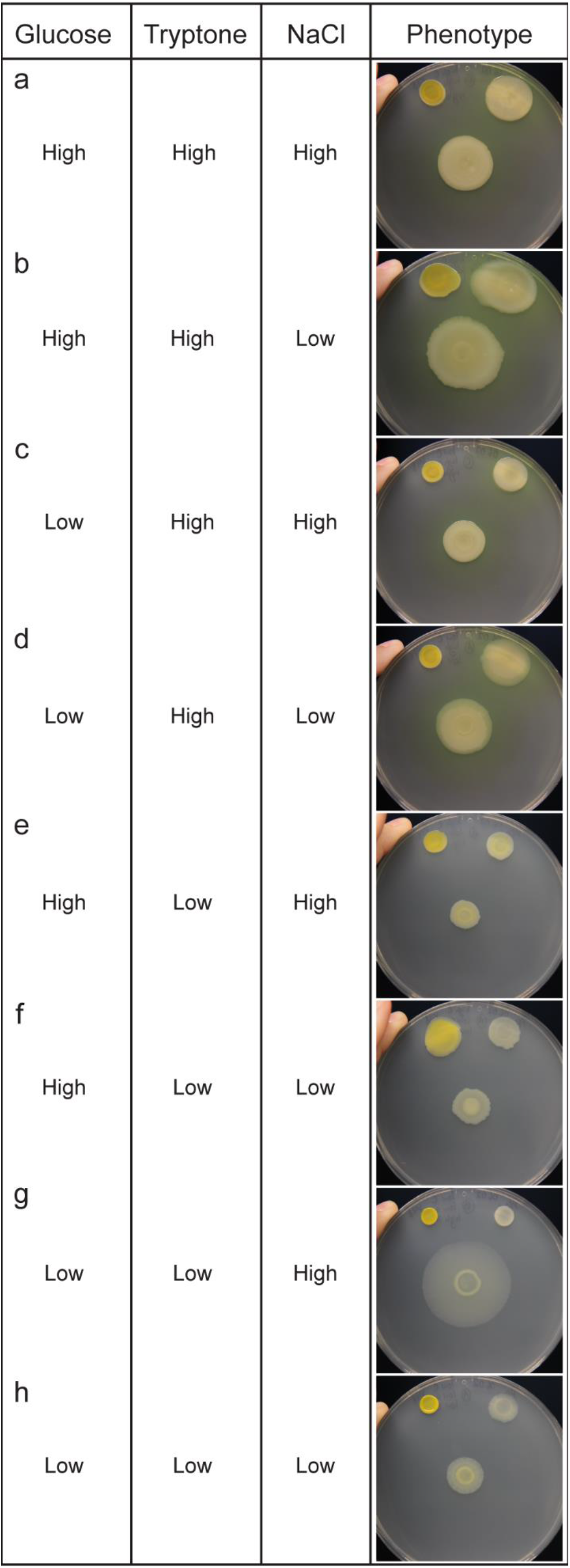
Effect of nutrient levels in interaction between *P. fluorescens* Pf0-1 and *Pedobacter* sp. V48, looking at 3 core components of TSB: tryptone (20 g/L), D-glucose (2.5 g/L), and NaCl (5 g/L) for ‘high’ concentrations. Components were reduced to 1/10 for ‘low’ concentrations. For all panels *Pedobacter* mono-culture is on the top left of the plate, *P. fluorescens* is on the top right of the plate, and the mixed colony is in the center. Pictures were taken 144 h after inoculation. Colonies were grown on 100 mm petri dishes.

Based on these results, we conclude that full interspecies social spreading was only observed in low nutrient medium supplemented with NaCl (Figs. 6b, d, and 7g). We observed reduced social spreading on low nutrient media without salt supplementation (Figs. 6c and 7h), and an absence of social behavior on rich media (Figs. 6a and 7a-f). While we can implicate salt as an important factor in social spreading, high salt concentrations alone are not sufficient to induce social behavior, as we do not see social behavior under rich media conditions. This indicates that there may be more than one important nutritional component factored in the decision of these bacteria to socialize.

## Discussion

In this study we investigate interspecies social spreading, a phenomenon that emerges from the interaction of two distantly-related soil bacteria. Neither species moves on its own under the conditions of our study, but a mixture of the two species can spread across a hard surface (2% agar). Contact between the two bacterial colonies is required for spreading to initiate, and this association is maintained as the co-culture expands. The social phenotype could be observed only under specific nutritional conditions, indicating an interplay between environmental and biological factors. The interaction between *Pedobacter* and *P. fluorescens* serves as a simple and tractable model for investigating interspecies interactions. Our research contributes to the growing body of work studying bacteria in social contexts to investigate emergent traits and behaviors.

Surface motility is a trait that could be beneficial to bacteria under a range of environmental conditions. Species related to *Pedobacter* sp. V48 use gliding motility on 1% agar or glass surfaces (27–29). V48 has not been observed to engage in gliding motility, but we have observed phenotypes similar to sliding motility in other species (30, 31), when inoculated on semi-solid agar (unpublished observations). *P. fluorescens* Pf0-1 is capable of flagella-driven swimming in and swarming motility on semi-solid agar (0.3% and 0.6% respectively) without the need for a partner bacterium (32, 33). Interspecies social spreading is distinct from *Pseudomonas* flagellar motility in its requirement of the presence of a second species. Additionally, media with higher agar percentages form environments that are non-permissive for flagella-driven motility in *P. fluorescens*, as well as most species, but together, Pf0-1 and V48 appear to employ an alternative strategy for movement across hard surfaces.

De Boer *et al*. (5) suggested that in water agar, the sporadic occurrence of movement they observed indicated a strategy by *Pedobacter* to escape competition. However, the co-migration under our conditions does not support this hypothesis, as the two species remain associated throughout the colony. Our contact experiments provide further evidence, as the presence of *Pedobacter* in the motile areas surrounding the *P. fluorescens* colony shows it has moved towards its partner, rather than away from it. The pattern of *Pedobacter* migration clearly indicates that it is not escaping.

Evidence, both from culturing and fluorescent imaging, shows that *P. fluorescens* and *Pedobacter* co-migrate across the hard agar surface. Initiation of the process requires physical contact, as motility is precluded when a semi-permeable membrane is placed between the two colonies. We suggest that the nature of this interaction is distinct from contact-dependent toxin delivery systems, such as type VI secretion and contact-dependent growth inhibition, as they commonly mediate signal exchange between closely-related species, and are involved in competition between more distantly-related strains (34–36). While our results do not rule out quorum sensing for communication between the two species (37), a diffusible signal (if it exists) does not appear be sufficient to trigger the motility response. Additionally, our experiment in which bacteria are pre-grown on cellophane indicates that social spreading is not triggered by a change in the medium caused by metabolic activity of one of the two species. Our data indicate that physical association is required for social spreading between *P. fluorescens* and *Pedobacter*. The question remains, are the bacteria producing a signal which induces an already-present motility mechanism in one species, or are they directly manipulating the environment in a way which facilitates co-migration, such as by production of a surfactant? Regardless of which mechanism is used, close association is still a prerequisite for either induction or facilitation of social spreading.

Bacteria dwelling in soil experience variations in a wide range of abiotic conditions, including the key parameters we have tested: salinity and available carbon and nitrogen (2). Environmental conditions have previously been shown to affect motility of individual species; gliding motility in some *Flavobacterium* species increases with reduced nutrient concentration (38, 39). Changes in behavior resulting from environmental fluctuations can affect how species interact with one another. The ability of *P. fluorescens* and *Pedobacter* to spread socially is dependent upon the conditions in which they are growing. In general, high concentrations of glucose and amino acids lead to a build-up of biomass and no apparent social movement.

Lower glucose and amino acid concentrations are associated with interspecies social spreading across the plate, but decreasing the salt concentration of the media slows expansion of the colony. Social spreading resulting from the interaction is conditional, with alteration of just a subset of environmental factors resulting in dramatic changes in behavior. It is tempting to speculate that the consortium of *P. fluorescens* and *Pedobacter* can integrate signals from each other’s presence and from the nutrient conditions of their environment to determine whether to behave socially. We see similar examples of intraspecies social behaviors being influenced both by biotic factors (quorum sensing) and by abiotic factors (nutrient conditions) in *P. aeruginosa* (40), *Bacillus subtilis* (41), and yeast (42).

There are a wide variety of examples of motility resulting from interspecies interactions, where the presence of a motile partner fosters the motility of an immotile participant. Non-motile *Staphylococcus aureus* hitchhikes on swimming *P. aeruginosa* (43) and *Burkholderia cepacia* co-swarms with *P. aeruginosa* in environments where it cannot do so independently (19). *X. perforans* induces motile *P. vortex* to swarm towards it, which allows it to hitchhike on top of *P. vortex* rafts (20). *P. vortex* is also capable of carrying fungal spores or antibiotic-degrading cargo bacteria to cross unfavorable environments (18, 44). In an even more complex system, *Dyella japonica* can migrate on fungal hyphae, but some strains can only do so in the presence of a *Burkholderia terrae* helper (45, 46). All of these examples of ‘hitchhiking’ phenomena require one species to already be motile, and stand in contrast to the behavior we have investigated, where social spreading emerges from two conditionally non-motile participants. The fact that both species are present at the edge of the spreading colony suggests that both have an active role in the behavior, though it doesn’t rule out the possibility of one species inducing motility in the other and hitchhiking, as seen in other systems (20).

In addition to describing a new mode of motility, this discovery highlights the possibility that many functions and behaviors of bacteria in complex communities may be triggered by interactions between different species or even domains. Studying interactions between two or more microorganisms may lead to the discovery of emergent traits that would be impossible to predict based on the study of each organism in isolation. Alongside approaches that characterize the members and connectedness of microbial communities, tools to decipher the phenotypic outcomes of interactions are needed in order to develop a full appreciation of microbiomes. Studies of this type are important for understanding the role of microbial communities within an ecological context.

We have investigated an interaction-dependent trait which emerges under particular nutritional conditions when distantly-related bacteria come into close physical contact. This interaction gives the participating bacteria the ability to spread on a hard agar surface, which neither can do alone. This strategy of co-migration may serve as an additional mechanism by which plant- and soil-associated bacteria can move in their natural environments, when the conditions do not favor the modes of single-species motility previously described. Given the distant and different locations from which these two strains were isolated, we hypothesize this is not a unique interaction between this pair, but rather has evolved between various *Pedobacter* and *Pseudomonas* species. To understand the phenomenon, several lines of investigation should be pursued: mechanistic studies which explore the factors each species is contributing to social spreading, the process by which contact triggers motility, whether there are important metabolic interactions, and the way in which environmental conditions are integrated into the decision to move together. The system we study is a tractable model for studying interspecies interactions, giving us the opportunity to answer questions about the nature of interspecies social spreading and ask questions about the broader field of bacterial communities. Models such as these will ultimately lead to a greater understanding of the functions of communities as a whole rather than as collections of individuals.

## Materials and Methods

### Bacterial strains, primers, plasmids, and culture conditions

Bacterial strains and plasmids are described in Table 1. *E. coli* was grown at 37 °C in LB Broth, Miller (Fisher Scientific). *Pseudomonas fluorescens* Pf0-1 and *Pedobacter* sp. V48 were routinely grown at 30°C or 20°C respectively, in 10% strength Tryptic Soy Broth (BD Difco™) amended with NaCl and KH_2_PO_4_, as described by de Boer (5). This medium is referred to throughout the text as TSB-NK. To differentiate the two species from mixed cultures we used *Pseudomonas* minimal medium (PMM) with 25 mM succinate (47) for *P. fluorescens* and 14.6 mM lactose for *Pedobacter*.

**Table 1.**
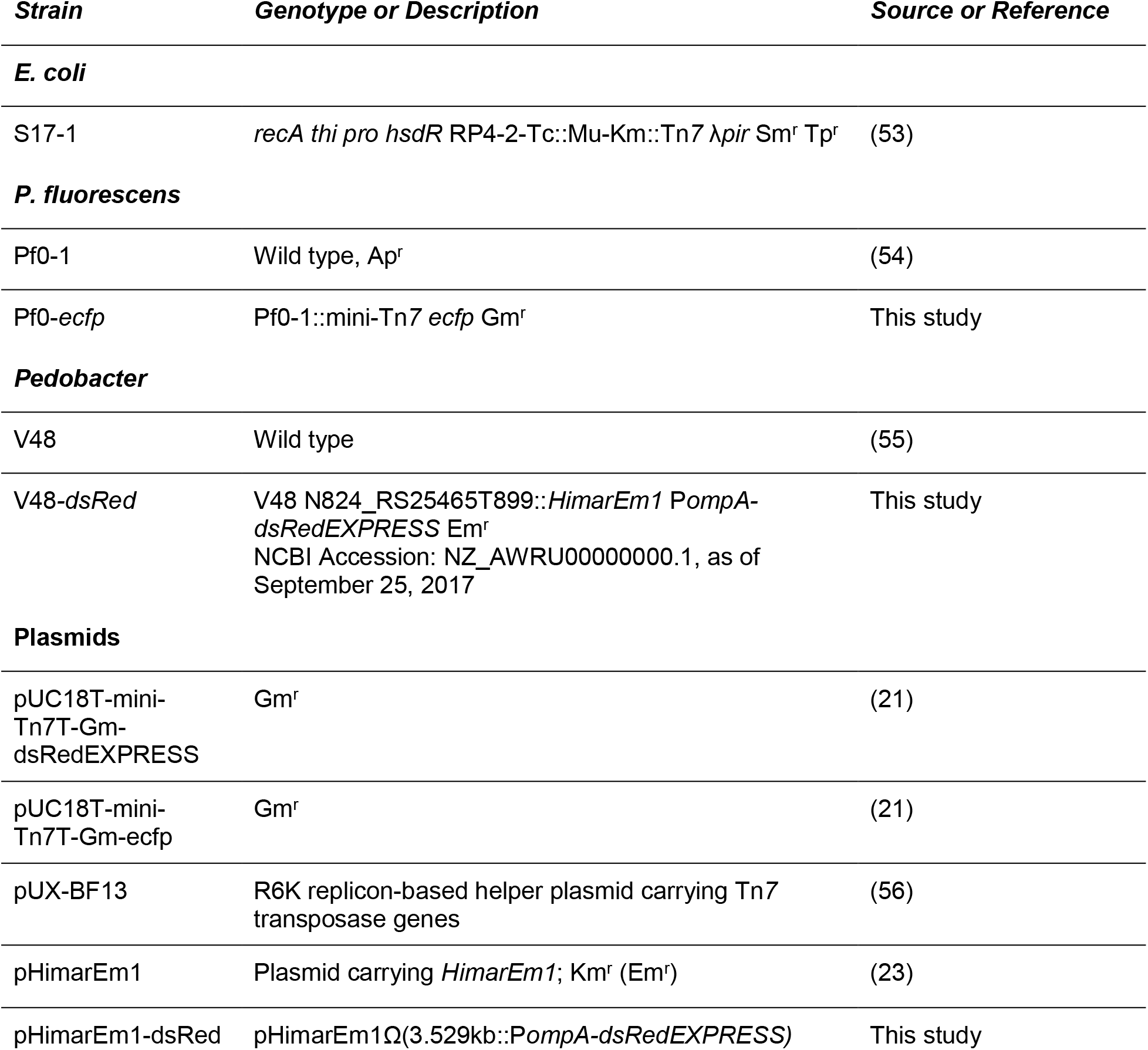
Bacterial strains and Plasmids.

Media were solidified with BD Difco™ Bacto™ agar (1.5% w/v) when required, except for social spreading assays, for which 2% agar was used. For experiments with variations in nutrients, we used full-strength TSB (30 g/L), 10% TSB (3 g/L), and 10% TSB amended with NaCl or KH_2_PO_4_ (called TSB-N or TSB-K, respectively), and a medium composed of D-glucose (2.5 g/L), tryptone (20 g/L), and NaCl (5 g/L). These individual components were used at those concentrations or reduced to 10% concentration in all eight combinations. For selection of transposon insertions carrying fluorescent protein genes, Kanamycin (50 μg/mL), Gentamicin (50 μg/mL), or Erythromycin (100 μg/mL) was added to the growth medium.

### Interspecies social spreading assays

*P. fluorescens* and *Pedobacter* for use in social spreading assays were incubated in 2 mL TSB-NK at 20 °C for 24 hours, with shaking (160 rpm). Social spreading assays were carried out on TSB-NK solidified with 2% agar. Plates were poured at a temperature of ~62 °C in a single layer and allowed to set for ~15 minutes before inoculation. Inoculation was done on freshly-poured plates.

i. Mixed inoculum assays. Assays were started by combining 5 μL of each participant in one spot on the agar surface. As controls, 10 μL spots of each bacterial isolate were plated distant from each other and the co-culture, all on the same plate. Once the inoculation liquid had dried, plates were incubated at 20 °C. Measurements of the colony diameter were taken every 24 hours. Experiments were performed in triplicate.
ii. Direct contact assay - adjacent plating. *P. fluorescens* and *Pedobacter* were grown as described above. The aliquots of bacteria were plated adjacent but without the drops touching. Once the inoculation liquid had dried, plates were incubated at 20 °C and monitored daily to determine the time at which colony growth led to contact between the isolates, and when spreading phenotypes developed.
iii. Direct contact assay - separation by membranes. *P. fluorescens* and *Pedobacter* were plated close together, separated only by a membrane. Either Millipore Polyethersulfone (PES) Express Plus^®^ Membrane (0.22 μm pores) or Gelman Sciences mixed-ester cellulose Metricel Membrane (0.45 μm pores) were cut into rectangular strips and sterilized by autoclaving. These strips were then embedded into the agar by suspending them perpendicular to the bottom of petri dishes with forceps, as agar was poured into plates. Once set, the filters protruded approximately 5 mm above the agar surface. Bacteria were inoculated on either side of the filter, with 5 μL spots of each species, close enough to touch the filter.
iv. Cellophane overlay assay. Squares of porous cellophane (GE Healthcare Bio-sciences Corp) were placed on top of TSB-NK plates. Cultures of *P. fluorescens, Pedobacter*, and a co-culture of the two, were placed on top of the cellophane, with cellophane alone used as a negative control. Plates were incubated at 20 °C for two days, at which point cellophane was removed, and 5 μL spots of either species were placed in the center of the plate, so that cultures were on a plate where cellophane had been (negative control), one where the partner species had been cultured, or one where a mix of the species had been cultured.
v. Heat-kill assay. Cells were scraped from TSB-NK plates, suspended in PBS buffer, heat-killed, and added on top of or adjacent to a colony of *Pseudomonas* or *Pedobacter* to test the ability of heat-killed cells to induce movement in the partner species. To place the heat-killed suspension adjacent to living colonies, a well was made in freshly-poured agar, by cutting a core using the top end of a 10 μL pipette tip (USA Scientific, Inc.), and partially filling it in using 60-70 μL agar. Cultures of *Pseudomonas* or *Pedobacter* (5 μL spots) were inoculated adjacent to the well, and the well was filled with the heat-killed suspension. For experiments in which the heat-killed suspension was added directly on top of living colonies, these colonies were initiated with 10 uL spots of liquid culture. The suspensions added directly on top of the colony or to the wells were heat-killed *Pseudomonas* or co-culture on/next to a *Pedobacter* colony, or heat-killed *Pedobacter* or co-culture on/next to a *Pseudomonas* colony. These heat-killed cells, or PBS buffer as a negative control, were added to the colonies or wells every 24 hours until the end of the experiment. The cells added on top of the colonies or into the wells were extracted from 4-day-old mono- and co-culture colonies on TSB-NK, inoculated and cultured as previously described. Whole colonies from these plates were resuspended in 1 mL PBS buffer, vortexed until fully suspended, then heat-killed at 65 °C for 15 minutes. Effectiveness of heat-killing was evaluated by plating 100 μL of resuspension on TSB-NK, and PMM with succinate or lactose.

### Fluorescent protein tagging

i. eCFP labeling of *P. fluorescens*. pUC18T-mini-Tn7T-Gm-ecfp was a gift from Herbert Schweizer (Addgene plasmid # 65030). A constitutively-expressed fluorescent protein gene carried by pUC18T-mini-Tn7T-Gm-ecfp was transferred to *P. fluorescens* by conjugation from *E. coli* S17-1, with transposase being provided by pUX-BF13 introduced from a second *E. coli* S17-1 donor, as previously described (48). Transposon-carrying strains were selected by growth on Gentamicin (50 μg/mL), and transposition of the miniTn7 element into the target site in the *P. fluorescens* genome was confirmed by PCR using primers Tn7-F and glmS-R (Table 2). Pf0-1 with fluorescent inserts were tested for alteration in interspecies social spreading by co-culturing with *Pedobacter*, as described above.
ii. dsRedEXPRESS labeling of *Pedobacter*. pUC18T-mini-Tn7T-Gm-dsRedExpress was a gift from Herbert Schweizer (Addgene plasmid #65032). To express *dsRedEXPRESS* in *Pedobacter*, a *Pedobacter* promoter was cloned upstream of the *dsRedEXPRESS* coding sequence. A highly expressed gene from an unpublished RNAseq experiment was identified (N824_RS25200) and the upstream 300 bp were amplified from *Pedobacter* genomic DNA using primers *PompA* and *dsRed*, designed for splicing-by-overlap extension-PCR (SOE-PCR) (Table 2). The promoter was then spliced with the amplified *dsRedEXPRESS* coding sequence using SOE-PCR (49). Flanking primers were designed with *KpnI* restriction sites, enabling cloning of the spliced product into a *KpnI* site in *pHimarEm1* (23). To join compatible ends between the plasmid and the amplicons, we used T4 DNA ligase (New England Biolabs, Inc.). The ligated plasmid was introduced into *E. coli* S17-1 competent cells by electroporation (BioRad Micropulser™). S17-1 colonies carrying the plasmid were selected by plating on LB medium containing Kanamycin (50 μg/mL), and the presence of the *dsRedEXPRESS* gene was confirmed by PCR, using *pHimar* KpnI-flank primers (Table 2). The resulting plasmid is called *pHimarEm1-dsRed*.

**Table 2.**
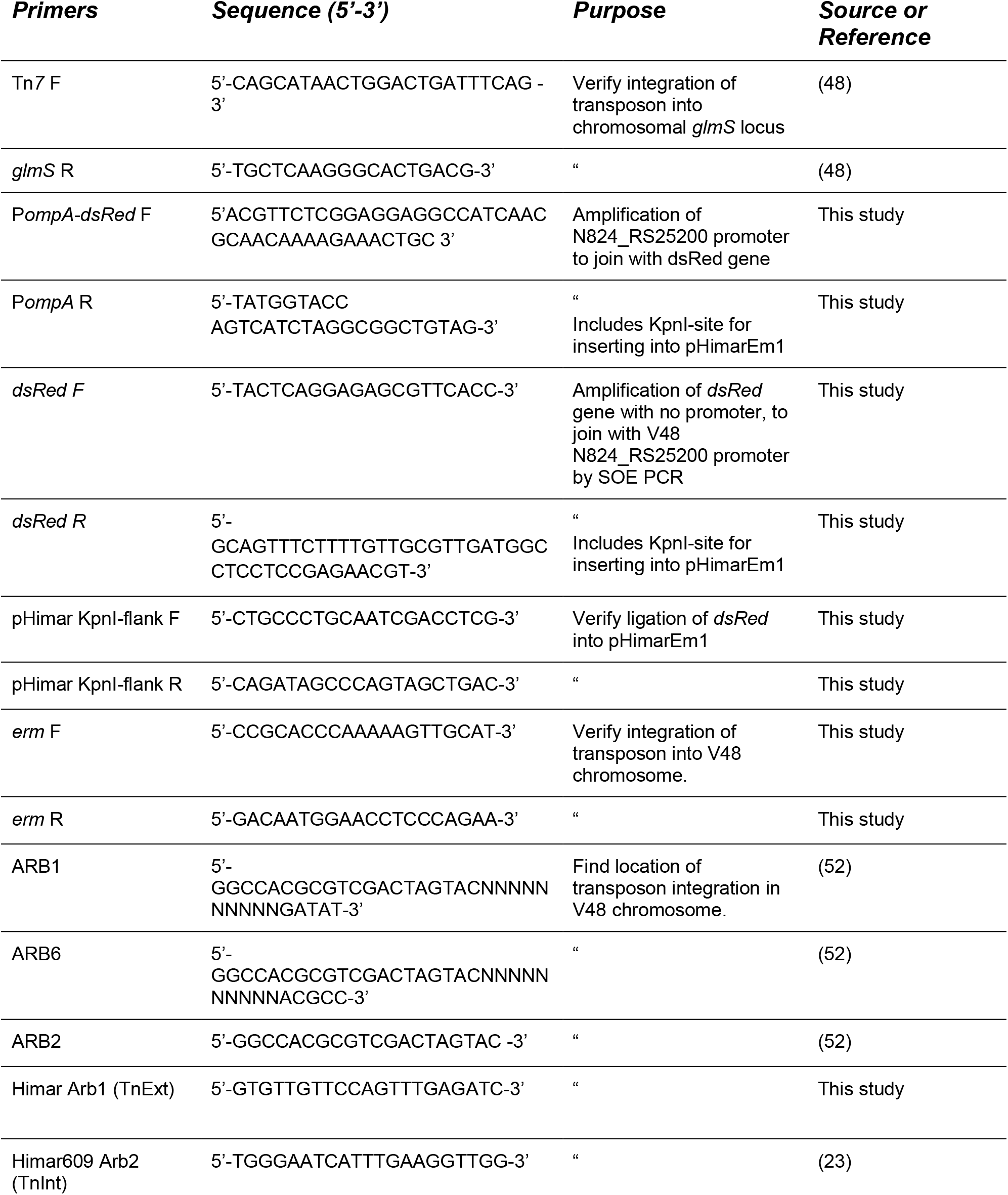
Primers.

*pHimarEm1-dsRed* was transferred to *Pedobacter* by conjugation using a method adapted from Hunnicutt and McBride, 2000. Briefly, 20 hour old cultures of *E. coli* S17-1 *(pHimarEm1-dsRed)* and *Pedobacter* were subcultured 1:100 into fresh LB, and grown to mid-exponential phase (*E. coli*) or for 7 hours *(Pedobacter)*. Cells were collected by centrifugation, suspended in 100 μL of LB, and then mixed in equal amounts on TSB-NK with 100 μL of 1M CaCl_2_ spread on the surface. Following overnight incubation at 30 °C, cells were scraped off the surface of the plate, and dilutions were plated on TSB-NK with Erythromycin (100 μg/mL) to select for strains that received the plasmid *(ermF* is not expressed in *E. coli*). Transconjugants were incubated at 25 °C for 3-4 days. Presence of the transposon insert in *Pedobacter* was confirmed using *ermF* primers (Table 2). *Pedobacter* with fluorescent inserts were tested for alteration in interspecies social spreading by co-culturing with *Pseudomonas*, as described above.

The transposon insertion sites in the *Pedobacter* chromosome were amplified by arbitrarily-primed PCR (51), using a method adapted from O’Toole *et al*. (52) (see table 2 for primers), and identified by sequencing the arb-PCR products. Nucleic acid sequencing was performed by Massachusetts General Hospital CCIB DNA Core. Sequences were analyzed using CLC Genomics Workbench Version 10.1.1 (QIAGEN) to find location of transposon integration.

### Imaging

Still pictures were taken using an EOS Rebel T3i camera (Canon) and processed using Photoshop CC 2017 Version: 14.2.1 and Illustrator CC 2017 Version: 17.1.0 (Adobe). Using Photoshop, the levels of some images were adjusted to improve contrast.

For microscopy, motile colonies were examined using an Axio Zoom.V16 microscope (Zeiss). To visualize fluorescent strains, filter set 43 HE DsRed was used with a 1.5 s exposure, shown with pseudo-color orange, as well as filter set 47 HE Cyan Fluorescent Protein, with a 600 ms exposure, shown with pseudo-color turquoise. Images were captured using Axiocam 503 mono camera, with a native resolution of 1936×1460 pixels. For image acquisition and processing we used Zen 2 Pro software (Zeiss).

### Statistics

We measured the amount of colony expansion of the mono-cultures of both *P. fluorescens* and *Pedobacter* and the expansion of social spreading in co-culture. Colony diameter of three independent experiments was measured every 24 hours. To compare the diameter of mono-cultures and co-cultures at each time point, we performed a two-way ANOVA followed by a Bonferroni post-hoc test, using GraphPad Prism version 5.04 for Windows (Graphpad Software).

We compared the movement speed between a combination of wild type *P. fluorescens* and *Pedobacter* to a combination of fluorescently-tagged Pf0-ecfp and V48-dsRed. Colony diameter of six independent experiments were measured every day, and speed was calculated by dividing the distance traveled by the amount of time elapsed since the last time point. To calculate average speed, we only used time points after the interspecies social spreading phenotype developed. To compare the means of the speed of the wild-type and tagged strains, we conducted an unpaired, two-tailed, Student’s t-test, using GraphPad Prism version 5.04.

## Acknowledgements

This research received no specific grant from any funding agency in the public, commercial, or not-for-profit sectors. LMM was supported by a University of Massachusetts Dartmouth Distinguished Doctoral Fellowship.

The authors would like to thank Brianna Arruda, Michael Baym, Jacob Palmer, Emma Piatelli, and Marian Wahl for constructive criticism and expert advice.

## Author contribution statement

LMM designed and carried out experiments, analyzed data, wrote the manuscript. ASB, SCS, and LMS each contributed a key experiment, and edited the manuscript. MWS contributed to experimental design, data analysis, writing and editing of the manuscript.

